# PerSVade: Personalized Structural Variation detection in your species of interest

**DOI:** 10.1101/2021.11.23.469703

**Authors:** Miquel Àngel Schikora-Tamarit, Toni Gabaldón

**Affiliations:** Barcelona Supercomputing Centre (BSC-CNS). Plaça Eusebi Güell, 1-3 08034 Barcelona, Spain; Institute for Research in Biomedicine (IRB Barcelona), The Barcelona Institute of Science and Technology, Baldiri Reixac, 10, 08028 Barcelona, Spain; Catalan Institution for Research and Advanced Studies (ICREA), Barcelona, Spain; Centro Investigación Biomédica En Red de Enfermedades Infecciosas, Barcelona, Spain

## Abstract

Structural variants (SVs) like translocations, deletions, and other rearrangements underlie genetic and phenotypic variation. SVs are often overlooked due to difficult detection from short-read sequencing. Most algorithms yield low recall on humans, but the performance in other organisms is unclear. Similarly, despite remarkable differences across species’ genomes, most approaches use parameters optimized for humans. To overcome this and enable species-tailored approaches, we developed perSVade (personalized Structural Variation Detection), a pipeline that identifies SVs in a way that is optimized for any input sample. Starting from short reads, perSVade uses simulations on the reference genome to choose the best SV calling parameters. The output includes the optimally-called SVs and the accuracy, useful to assess the confidence in the results. In addition, perSVade can call small variants and copy-number variations. In summary, perSVade automatically identifies several types of genomic variation from short reads using sample-optimized parameters. We validated that perSVade increases the SV calling accuracy on simulated variants for six diverse eukaryotes, and on datasets of validated human variants. Importantly, we found no universal set of “optimal” parameters, which underscores the need for species-specific parameter optimization. PerSVade will improve our understanding about the role of SVs in non-human organisms.

## INTRODUCTION

Structural variants (SVs) are large changes (typically >50 bp) in the DNA between individuals that alter genome size (duplications and deletions) or generate rearrangements (inversions, translocations and interspersed insertions) (*1*, *2*). In eukaryotes, SVs can drive clinically-relevant phenotypes including cancer (*3*–*5*), neurological diseases (*6*, *7*) or antifungal drug resistance (*8*, *9*). In addition, SVs may generate significant intraspecific genetic variation across many taxa like humans (*10*–*12*), songbirds (*13*) or rice plants (*14*). Despite their role on human health and natural diversity, most genomic studies overlook SVs due to technical difficulties in calling SVs from short reads (*15*). This means that the role of SVs remains largely unexplored across eukaryotes.

Inferring SVs from short reads is challenging because it relies mostly on indirect evidence coming from *de novo* assembly alignment, changes in read depth or the presence of discordantly paired / split reads in read mapping analysis (*16*–*21*). Long-read based SV calling may avoid some of these limitations, but short read-based SV calling remains a cost-effective strategy to find SVs in large cohorts (*14*, *15*, *22*). Recent benchmarking studies compared the performance of different tools in human genomes and found that SV calling accuracy is highly dependent on the methods and filtering strategy used (*15*, *23*, *24*). Such studies are useful to define ‘best practices’ (optimal methods and filtering strategies) for SV calling in human samples. However, few studies have investigated the accuracy of these tools on non-human genomes. It is unclear whether the human-derived ‘best practices’ for SV calling can be reliably used in other species. We hypothesize that this may not be the case for genomes with different contents of repetitive or transposable elements, which constrain the short read-based SV calling accuracy (*24*). In summary, current tools for short-read based SV calling are often unprepared for non-human genomes, which hinders the study of SVs in most organisms.

To overcome this limitation, we developed the *personalized Structural Variation detection* pipeline, or perSVade (pronounced “persuade”), which is designed to adapt a state-of-the-art SV calling pipeline to any genome/species of interest. PerSVade detects breakpoints (two joined regions that exist in the sample of interest and not in the reference genome) from short paired-end reads and summarizes them into complex SVs (deletions, inversions, tandem duplications, translocations and interspersed insertions). The pipeline provides automated benchmarking and parameter selection for these methods in any genome or sequencing run, which is useful for species without such recommended parameters. PerSVade provides an automated report of the SV calling accuracy on these simulations, which serves to estimate the confidence of the results on real samples. Beyond SV detection, perSVade can be used to find small variants (Single Nucleotide Polymorphisms (SNPs) and insertions/deletions (IN/DELs)) and read depth-based Copy Number Variation (CNV), all implemented within a flexible and modular framework.

The following sections describe perSVade and its SV calling performance on various datasets of both simulated and real genomes with SVs.

## MATERIALS AND METHODS

### PerSVade pipeline

PerSVade has several modules that can be executed independently (each with a single command) and/or combined to obtain different types of variant calls and functional annotations. The following sections describe how each of these modules work, and **Figure 1** shows how they can be combined.

**Figure 1.**
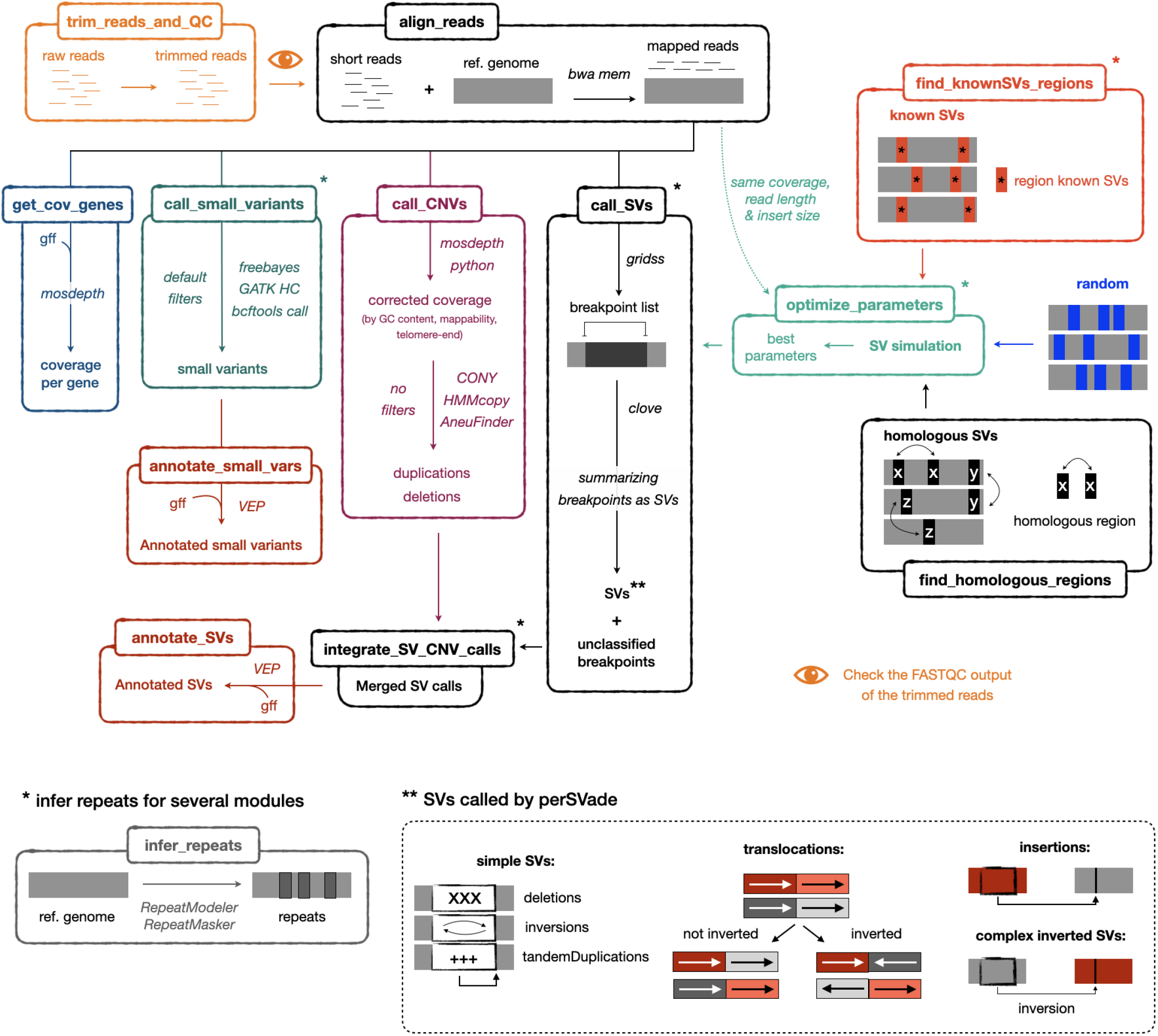
Schematic representation of the modular workflow of PerSVade. This figure shows the modules of perSVade (each represented in a different box and executable with a single command), which may be combined following the drawn arrows. The italic text describes the algorithms used at each step. The pipeline identifies either structural variants (SVs) (module ‘call_SVs’), coverage-derived copy number variants (CNVs) (module ‘call_CNVs’), small variants (module ‘call_small_variants’) and/or changes in the coverage per gene (module ‘get_cov_genes’) from aligned short paired-end reads (obtained with the module ‘align_reads’). The different types of SVs output by ‘call_SVs’ are drawn at the bottom for clarity. In addition, the module ‘trim_treads_and_QC’ can be used to trim the reads and perform quality control with *FASTQC* before read alignment. On another note, several modules (‘call_SVs’, ‘find_knownSVs_regions’, ‘integrate_SV_CNV_calls’, ‘optimize_parameters’ and ‘call_small_variants’) use an annotation of genomic repeats that can be obtained with the module ‘infer_repeats’ (bottom left). The most novel aspect of perSVade is the automatic parameter optimization for SV calling adapted to the input (implemented in the module ‘optimize_parameters’). This is achieved through simulations of SVs on the reference genome, which can be randomly placed (‘random’), around regions with previously known SVs (‘known’) or on regions with pairwise homology (‘homologous’). The modules ‘find_knownSVs_regions’ and ‘find_homologous_regions’ can be used to infer these ‘known’ and ‘homologous’ regions, respectively. In addition, the variants found with ‘call_SVs’ and ‘call_CNVs’ can be combined with the module ‘integrate_SV_CNV_calls’. Finally, the modules ‘annotate_SVs’ and ‘annotate_small_vars’ can be used to obtain a functional annotation of the variants. See **Materials and Methods** for more details. In addition, note that **Figure S1** includes a more detailed representation of how ‘optimize_parameters’ works.

#### Module ‘trim_reads_and_QC’

This module runs *trimmomatic* (*25*) (v0.38) with default parameters for the input reads followed by *fastqc* (https://www.bioinformatics.babraham.ac.uk/projects/fastqc, v0.11.9) on the trimmed reads. These trimmed reads may be used for downstream analysis after checking that they are reliable according to the output of *fastqc*.

#### Module ‘align_reads’

This module runs *bwa mem* (http://bio-bwa.sourceforge.net/bwa.shtml, v0.7.17) to align the short reads, generating a sorted .bam file (using *samtools* (*26*) (v1.9)) with marked duplicates (through *GATK MarkDuplicatesSpark* (https://gatk.broadinstitute.org/hc/en-us/articles/360036358972-MarkDuplicatesSpark, v4.1.2.0)), that is the core input of several downstream modules (‘call_SVs’, ‘optimize_parameters’, ‘call_CNVs’, ‘call_small_variants’ and ‘get_cov_genes’).

#### Module ‘call_SVs’

This module uses *gridss (21, 27)* to infer a list of breakpoints (two regions of the genome -two breakends-that are joined in the sample of interest and not in the reference genome) from discordant read pairs, split reads and *de novo* assembly signatures. The breakpoints are summarized into SVs with *clove* (*28*) (v 0.7). Importantly, this module (and others) runs *clove* without the default coverage filter to classify deletion-like (DEL-like) and tandem duplication-like (TAN-like) breakpoints into actual deletions and tandem duplications. Instead, perSVade ‘call_SVs’ calculates the relative coverage of the regions spanned by such breakpoints (using *mosdepth* (*29*)). This information is used to define the final set of deletions (DEL-like breakpoints with a coverage below a “max_rel_coverage_to_consider_del” threshold) and tandem duplications (TAN-like breakpoints with a coverage above a “min_rel_coverage_to_consider_dup” threshold). This setting allows a separate thresholding for the classification of DEL and TAN-like breakpoints, which is a novel feature of perSVade as compared to the current implementation of *clove*. Note that this module requires as an input a set of parameters to filter the *gridss* and *clove* outputs, which may be inferred using the module ‘optimize_parameters’ (described below).

The final output of this module is a set of files with the called variants, which belong to these types:

- Simple SVs: deletions, inversions and tandem duplications (duplication of a region which gets inserted next to the affected region). This module outputs one .tab file for each of these SV types.
- Translocations: whole-arm balanced translocations between two chromosomes, which can be inverted or not. There is one .tab file for translocations.
- Insertions: a region of the genome is copied or cut and inserted into another region. Note that these are not *de novo* insertions (i.e. of DNA not present in the reference), which are actually not called by perSVade. There is one .tab file for insertions.
- Unclassified SVs: One .tab file reports all the variants that are called by *clove* and cannot be assigned to any of the above SV types. These include *clove*’s unclassified breakpoints (which could be part of unresolved/unknown complex variants) and complex inverted SVs (which are non-standard SVs). These types of SVs are not included in the simulations performed by ‘optimized parameters’ (see below), so that their accuracy is unknown. This is why we group them together into a single file.

#### Module ‘optimize_parameters’

To find optimal parameters for running ‘call_SVs’ in a given input dataset, this module generates two simulated genomes with up to 50 SVs of each of five types (insertions, translocations, deletions, inversion and tandem duplications) with *RSVsim (30)* (v1.28) and custom python (v3.6) scripts (which use *biopython (31)* (v1.73)). For each genome, the module simulates reads with *wgsim* (https://github.com/lh3/wgsim, v1.0) and *seqtk* (https://arc.vt.edu/userguide/seqtk/, v1.3) with a read length, insert size and coverage matching that of the input dataset. Note that the read simulation is performed according to a user-defined zygosity and ploidy (through the argument ‘*--simulation_ploidies*’) to resemble various organisms. For example, if ‘*--simulation_ploidies diploid_hetero*’ is specified, this module simulates heterozygous SVs by merging reads from both the reference genome and the simulated genome with SVs in a 1:1 manner. PerSVade ‘optimize_parameters’ then tries several combinations (>13,000,000,000 by default, although this can be user-defined) of parameters to run *gridss* and *clove* and filter their outputs. The detailed explanation about the used filters can be found in **Additional Materials and Methods.** One of these possible filters includes removing SVs that overlap repetitive elements, which may be inferred with the module ‘infer_repeats’ (see below). This module selects the combination of filters that yield the highest F-value (the harmonic mean between precision and recall) for each simulated genome and SV type (see **Additional Materials and Methods** for more information on how accuracy is calculated). These filters are optimised for each simulation, and thus may not be accurate on independent sets of SVs (due to overfitting). In order to reduce this effect, perSVade ‘optimize_parameters’ selects a final set of “best parameters” that work well for all simulations and SV types. This set of best parameters may be used in the ‘call_SVs’ module. The accuracy (F-value, precision, recall) of these parameters on each simulation and SV type is reported in a tabular file, which serves to evaluate the expected calling accuracy. All plots are generated using *python* (v3.6) and the libraries *seaborn* (https://seaborn.pydata.org/, v0.9.0) and *matplotlib* (https://matplotlib.org/, v3.3.0). In addition, the *python* packages *scipy* (*32*) (v1.4.1), *scikit-learn (33)* (v0.21.3), *psutil* (https://github.com/giampaolo/psutil, v5.7.2) and *pandas* (https://pandas.pydata.org/, v0.24.2) are used for scripting and various statistical calculations. On another line, *pigz* (https://zlib.net/pigz/, v2.4) and *gztool* (https://github.com/circulosmeos/gztool, v0.11.5) are used for fast compression steps. Finally, perSVade ‘optimize_parameters’ uses *picard* (http://broadinstitute.github.io/picard/, v2.18.26) to construct a sequence dictionary for the provided reference genome.

By default, the simulated events are placed randomly across the genome. However, real SVs often appear around repetitive elements or regions of the genome with high similarity (e.g.: transposable elements insertions) (*24*, *34*–*36*). This means that random simulations may not be realistic, potentially leading to overestimated calling accuracy and a parameter selection unfit for real SVs (*24*). To circumvent this, perSVade ‘optimize_parameters’ can generate more realistic simulations occurring around some user-defined regions (i.e. with previously known SVs or homologous regions) provided with the --*regions_SVsimulations* argument. Importantly, perSVade provides an automatic way to infer such regions through the modules ‘find_knownSVs_regions’ and ‘find_homologous_regions’ (described below) In addition, note that **Figure S1** includes a detailed graphical representation of how this module works.

#### Module ‘find_knownSVs_regions’

This module finds regions with known SVs using a provided list of sequencing datasets (with the option *--close_shortReads_table*) from species close to the reference genome. These datasets are processed with perSVade’s modules ‘trim_reads_and_QC’, ‘align_reads’ and ‘call_SVs’ (using default parameters) to find SVs. This module then outputs a .bedpe file with the +-100bp regions around the breakends from these SVs. This .bedpe file can be input to the module ‘optimize_parameters’ through the *--regions_SVsimulations* argument in order to perform ‘known’ realistic simulations.

#### Module ‘find_homologous_regions’

This module infers homologous regions by defining genomic windows (from the reference genome) of 500 bp as a query for a *blastn (37)* against the same reference genome. Hits with an e-value <10^−5^ that cover >50% of the query regions are defined as pairs of homologous regions, which are written as a .bedpe file. This .bedpe file can be input to the module ‘optimize_parameters’ through the *--regions_SVsimulations* argument in order to perform ‘homologous’ realistic simulations.

#### Module ‘call_CNVs’

Copy Number Variants (CNVs) are a type of SVs in which the genomic content varies (deletions or duplications). The ‘call_SVs’ module (see previous section) identifies some CNVs (insertions, tandem duplications, deletions and complex inverted SVs) but it can miss others (i.e.: whole-chromosome duplications or regions with unknown types of rearrangements yielding CNVs (*8*, *38*)). PerSVade uses this ‘call_CNVs’ module to call CNVs from read-depth alterations. For example, regions with 0x or 2x read-depth as compared to the mean of the genome can be called deletions or duplications, respectively. A straightforward implementation of this concept to find CNVs is challenging because many genomic features drive variability in read depth independently of CNV (*39*, *40*). In order to solve this, perSVade ‘call_CNVs’ calculates the relative coverage for windows of the genome (using *bedtools (41)mosdepth (29)* (v0.2.6)) and corrects the effect of the GC content, mappability (calculated with *genmap (42)* (v1.3.0)) and distance to the telomere (using *cylowess* for nonparametric regression (https://github.com/livingsocial/cylowess) as in (*40*)). Note that *cylowess* uses the library *cython* (*43*) (v0.29.21). This corrected coverage is used by CONY (*44*) (v1.0), AneuFinder (*45*) (v1.18.0) and/or HMMcopy (*46*) (v1.32.0) to call CNVs across the genome. Note that we modified the R code of CONY to be compatible with the input corrected coverage. The corrected code (used in the pipeline) is available in https://github.com/Gabaldonlab/perSVade/blob/master/scripts/CONY_package_debugged.R. PerSVade ‘call_CNVs’ generates consensus CNV calls from the three programs taking always the most conservative copy number for each bin of the genome. For example, if the used programs disagree on the copy number of a region the closest to 1 will be taken as the best estimate.

#### Module ‘integrate_SV_CNV_calls’

This module generates a vcf file showing how SVs (called by the modules ‘call_SVs’ and/or ‘call_CNVs’) alter specific genomic regions. It also removes redundant calls between the CNVs identified with ‘call_SVs*’*and those derived from ‘call_CNVs’ (using *bedmap* from the *bedops* tool (*47*) (v2.4.39)). This is useful for further functional annotation. Each SV can be split across multiple rows when it affects more than one region of the genome. All rows related to the same SV are identified by the field variantID in INFO. On top of this, each row has a unique identifier indicated by the field ID. Some SVs generate de novo-inserted sequences around the breakends, and each of these is represented in a single row. Note that each of the rows may indicate a region under CNV (with the SVTYPE in INFO as DEL, DUP or TDUP), a region with some rearrangement (with the SVTYPE in INFO as BND) or a region with a de novo insertion (with the SVTYPE in INFO as insertionBND). See https://github.com/Gabaldonlab/perSVade/wiki/8.-FAQs#what-is-in-sv_and_cnv_variant_callingvcf for more information about the format of this .vcf file. We designed this vcf to include only these three types of regions because they can be interpreted by the Ensembl Variant Effect Predictor (*48*) for functional annotation.

#### Module ‘annotate_SVs’

This module runs the Ensembl Variant Effect Predictor (*48*) (v100.2) on the vcf output of the module ‘integrate_SV_CNV_calls’ to get the functional annotation of each SV. This requires a .gff file from the user.

#### Module ‘call_small_variants’

This module performs small variant (SNPs and small IN/DELs) calling with either *freebayes (49)* (v1.3.1), *GATK HaplotypeCaller* (*50*) (v4.1.2.0) and/or *bcftools call* (https://github.com/samtools/bcftools, v1.9) and integrates the results into .tab and .vcf files. **Additional Materials and Methods** provides further information on how this calling is performed.

#### Module ‘annotate_small_vars’

This module runs the Ensembl Variant Effect Predictor (*48*) (v100.2) on the vcf output of the module ‘call_small_variants’ to obtain the functional annotation of each variant. This requires a .gff file from the user.

#### Module ‘get_cov_genes’

This module runs *mosdepth (29)* (v0.2.6) *t*o obtain the coverage for each gene, which requires a .gff file from the user.

#### Module ‘infer_repeats’

This module annotates repetitive elements in a genome, which can be used for the modules ‘call_SVs’, ‘find_knownSVs_regions’, ‘integrate_SV_CNV_calls’, ‘optimize_parameters’ and ‘call_small_variants’.

These repeats are inferred with RepeatModeler (*51*) (v2.0.1) and RepeatMasker (*52*) (v4.0.9). The user can input these repeats to several modules (with --repeats_file), which will have the following effects:

- If repeats are provided, ‘optimize_parameters’ will assess whether removing SV calls overlapping repeats increases the overall accuracy. If so, the resulting optimized parameters will include a ‘filter_overlappingRepeats: True’. If you use these optimized parameters in ‘call_SVs’, any breakpoint overlapping repeats will be removed.
- If repeats are provided, ‘call_SVs’ may filter out SVs that overlap repeats if the SV filtering parameters include a ‘filter_overlappingRepeats: True’.
- If repeats are provided, ‘find_known_SVs’ will pass them to the ‘call_SVs’ module.
- If repeats are provided, ‘integrate_SV_CNV_calls’ will add a field in the INFO which indicates whether the SVs overlap repeats.
- If repeats are provided, ‘call_small_variants’ will add a field in the tabular variant calling file which indicates whether the SVs overlap repeats.

Alternatively, the user can specify ‘--repeats_file skip’ to avoid the consideration of repeats in all these modules.

### Testing SV calling with perSVade on simulated structural variants

To test perSVade’s performance on different species we ran it on paired-end WGS datasets for six eukaryotes (*Candida glabrata*, *Candida albicans*, *Cryptococcus neoformans, Arabidopsis thaliana*, *Drosophila melanogaster* and *Homo sapiens*). To obtain a high number of SVs we gathered three samples for each species with enough genetic divergence to the reference genome. For this, we first used an automatic pipeline to find these samples running the script https://github.com/Gabaldonlab/perSVade/blob/master/scripts/perSVade.py with the options *--close_shortReads_table auto --n_close_samples 3 --nruns_per_sample 1 --target_taxID <species_taxID>*. This used *entrez-direct* (https://www.ncbi.nlm.nih.gov/books/NBK179288/, v13.3), *SRA Tools* (https://github.com/ncbi/sra-tools, v2.10.9) and *ete3 (53)* (v3.1.2) to query the SRA database (*54*) and find three WGS datasets of close taxIDs (to each *<species_taxID>* according to the NCBI taxonomy species tree (*55*)) with a coverage >30x and >40% of mapped reads to the reference genome. We could find three such datasets for *C. albicans*, *C. neoformans, A. thaliana* and *D. melanogaster*, which included samples from the same species or genera as the target species, with >65% of the reads mapping to the reference genome. We randomly downsampled the *A. thaliana* and *D. melanogaster* runs to 30x coverage (using *samtools* (*26*) (v1.9)) for faster computation (using the option *--max_coverage_sra_reads 30*). For *C. glabrata* we used datasets generated in our lab from three divergent strains (BG2, CST34 and M12, from (*9*)). All these datasets are listed in **Table S1.** Finally, we tested perSVade on three *H. sapiens* datasets previously used for benchmarking SV callers (*23*, *24*). These included NA12878 (a Genome in a Bottle (GIAB) cell line related to the Ceph family (*56*, *57*)), HG002 (another GIAB project with reads available at ftp://ftp-trace.ncbi.nlm.nih.gov/giab/ftp/data/AshkenazimTrio/HG002_NA24385_son/NIST_HiSeq_HG002_Homogeneity-10953946/NHGRI_Illumina300X_AJtrio_novoalign_bams/HG002.hs37d5.60X.1.bam) and CHM1/CHM13 (two haploid cell lines sequenced independently (*58*), for which we merged the raw reads to generate synthetic diploid data). The reference genomes were taken from the Candida Genome Database (*59*) (version s02-m07-r35 for *C. glabrata* and ‘haplotype A’ from version A22-s07-m01-r110 for *C. albicans*), GenBank (*60*) (accession GCA_000149245.3 for *C. neoformans*, GCA_000001735.2 for *A. thaliana* and GCA_000001215.4 for *D. melanogaster*) and UCSC (*61*) (the latest version of genome hg38 at 06/04/2021 for *H. sapiens*, keeping only chromosomes 1-22, X,Y and the mitochondrial DNA). In addition, we performed quality control of the reads with fastqc (https://www.bioinformatics.babraham.ac.uk/projects/fastqc, v0.11.9) and trimming with *trimmomatic (25)* (v0.38).

We ran the SV calling pipeline of perSVade (using the modules ‘align_reads’, ‘call_SVs’ and ‘integrate_SV_CNV_calls’) on all these datasets using either ‘default’ or optimized parameters (based on ‘random’, ‘known’ or ‘homologous’ simulations using the modules ‘optimize_parameters’, ‘find_homologous_regions’ and ‘find_knownSVs_regions’). Note that we used the module ‘infer_repeats’ to find repetitive elements in each genome. These were provided to ‘optimize_parameters’ to assess whether filtering out repeats improved SV calling accuracy. In addition, we simulated diploid heterozygous SVs for the diploid genomes (*C. albicans*, *A. thaliana, D. melanogaster* and *H. sapiens*) and haploid SVs for the haploid genomes (*C. glabrata, C. neoformans*). We used computational nodes in an LSF cluster (https://www.ibm.com/support/pages/what-lsf-cluster) with 16 cores and either 32 Gb (for *C. glabrata*, *C. albicans*, *C. neoformans*), 64 Gb (for *A. thaliana* and *D. melanogaster*) and 96 Gb (for *H. sapiens*) of RAM for the testing. We first ran the read alignment step (module ‘align_reads’) for all samples, and then used the resulting .bam files as inputs for the other perSVade modules. We calculated the resource consumption (running time and maximum RAM used) for each of these perSVade runs, thus ignoring the resources related to read alignment. Of note, perSVade was run with different parameters for the human datasets to avoid excessive resource consumption and match our computational infrastructure. First, we skipped the marking of duplicate reads on the .bam files (default behavior) with perSVade’s *--skip_marking_duplicates* option on the module ‘align_reads’. Second, we ran the simulations on a subset of the genome (only chromosomes 2, 7, 9, X, Y and mitochondrial). Third, we skipped the ‘homologous’ simulations in human samples because we could not finish the inference of pairs of homologous regions (see previous section) due to excessive memory consumption. By running this inference on a few chromosomes we realised that there are millions of such regions, generating excessively large files. Finally, we tested the accuracy of all the optimized parameters (for each sample / simulation) on the other samples / simulations using the script https://github.com/Gabaldonlab/perSVade/blob/master/testing/get_accuracy_parameters_on_sorted_bam.py. **Additional Materials and Methods** provides further information on how accuracy is calculated.

### Testing perSVade on real SVs

To validate the usage of perSVade on real data we focused on public datasets with available short-reads and independently-defined sets of known SVs. We could find such SVs in the human samples (also used in the testing mentioned above), for which SV callsets of deletions or inversions exist (as done in (*24*)). We defined as ‘true SVs’ the deletions of NA12878 (defined in (*57*), available at ftp://ftp-trace.ncbi.nlm.nih.gov/giab/ftp/technical/svclassify_Manuscript/Supplementary_Information/Personalis_1000_Genomes_deduplicated_deletions.bed), the high-confidence deletions of HG002 (available at ftp://ftp-trace.ncbi.nlm.nih.gov/giab/ftp/data/AshkenazimTrio/analysis/NIST_SVs_Integration_v0.6/HG002_SVs_Tier1_v0.6.vcf.gz) and the union of all deletions and inversions found in either CHM1 or CHM13 lines (defined by (*58*), available at http://eichlerlab.gs.washington.edu/publications/Huddleston2016/structural_variants/).

We then tested the accuracy of the ‘training’ parameters optimized for each sample and simulation of the six eukaryotes mentioned above (in the section ‘Testing SV calling with perSVade on simulated structural variants’) on these human samples using the script https://github.com/Gabaldonlab/perSVade/blob/master/testing/get_accuracy_parameters_on_sorted_bam.py. In addition, we removed SVs overlapping simple repeats or low complexity regions (as inferred by the module ‘infer_repeats’) from this analysis. Note that each of these ‘true SV’ datasets were defined on different reference genomes: the NA12878 and HG002 callsets were based on hg19 and the CHM1/CHM13 was relative to hg38. This means that we could not directly use the optimized training parameters from the human samples from the previous section, since they were all based on hg38. We thus ran perSVade’s SV calling and parameter optimization modules on NA12878 and HG002 using the hg19 reference, and used the resulting optimum parameters as ‘training’ for these two samples. For this, we obtained the latest version of hg19 and hg38 genomes at 06/04/2021 from UCSC (*61*), keeping only chromosomes 1-22, X,Y and the mitochondrial DNA.

## RESULTS

### PerSVade: a pipeline to call and interpret structural variants in your species of interest

PerSVade identifies SVs from a paired-end WGS dataset and a reference genome as sole inputs. It identifies breakpoints from the aligned reads with *gridss (21)*, and summarizes them into actual SVs (insertions, translocations, deletions, inversion and tandem duplications) with *clove* (*28*). We followed the recent recommendation of using a single, high-performing algorithm for breakpoint calling instead of using multiple software (*24*). We chose *gridss* because of its high accuracy in several benchmarking studies (*23*, *24*). In addition, our pipeline generates a functional annotation of the variants, which is useful to evaluate the altered genomic regions and aid downstream analyses. In summary, perSVade is a pipeline to find and interpret SVs from most eukaryotic sequencing datasets **(Figure 1)**.

A key feature of perSVade is the parameter optimization step (implemented in ‘optimize_parameters’ module and shown in **Figure S1**). There are no specific recommendations for filtering the outputs of *gridss* and *clove* in most species, and it is unclear whether the parameters validated on model organisms are universal. Similarly, the performance of these algorithms on different sequencing formats (i.e. varying read lengths, coverage or insert size) is not easy to predict. To solve this automatically, perSVade ‘optimize_parameters’ generates simulated genomes (based on the reference genome and input dataset) with SVs and choses the most accurate filters (with the highest F-value) for these simulations. To account for different mechanisms of SV formation, the simulations can be either 1) randomly placed across the genome (“random” simulations), 2) around regions with previously known SVs (“known” simulations) or 3) around regions with homologous sequences (“homologous” simulations). We consider that “known” and “homologous” simulations are more realistic than the “random” ones. See **Materials and Methods** for further details. Regardless of the simulation type, the optimised filters can be used for the SV calling on real data, potentially yielding the highest possible performance. The accuracy of the optimised filters on different simulations is reported as a tabular file, which is useful to define the expected calling accuracy. We hypothesize that this accuracy may vary across species and/or sequencing formats, and perSVade can infer it on any input sample. All in all, perSVade automatically finds the best filters and reports the expected calling accuracy for each input sample.

We validated the usability of perSVade by running it on available sequences for six phylogenetically diverse eukaryotes with different genome sizes (*Candida glabrata* (12 Mb)*, Candida albicans* (14 Mb)*, Cryptococcus neoformans* (19 Mb)*, Arabidopsis thaliana* (120 Mb)*, Drosophila melanogaster* (144 Mb) and *Homo sapiens* (3163 Mb)), with three WGS runs per species (yielding datasets with 6,75·10^6^ - 1,59·10^9^ reads, **see Materials and Methods**). We ran the pipeline using parameter optimization with “random”, “known” or “homologous” simulations. In addition, we ran perSVade with default parameters as a baseline, useful to evaluate the impact of parameter optimization (the core and most novel feature of perSVade) on calling accuracy and resource consumption. We found that the computational burden (running time and memory used) was highly variable among datasets and correlated with genome and dataset sizes. As expected, parameter optimization increased resource consumption in all cases. This burden was particularly high for the human datasets, which may hinder the usage of perSVde on such large genomes **(Figure S2)**. However, we consider that such choices should be left to the user based on these results. Taken together, our analysis indicates that perSVade can be used for SV calling in a wide range of eukaryotes and sequencing datasets.

### PerSVade’s parameter optimization improves calling accuracy in simulated datasets

In order to clarify the impact of parameter optimization on calling accuracy we measured the performance of perSVade’s SV calling on these samples and simulations. We found that the F-value after parameter optimization on ‘random’ and ‘known’ simulations was high (between 0.75 - 1.0) in most samples and SV types (with one exception in *Drosophila melanogaster* that yielded an F-value ~ 0.5). The F-value on ‘homologous’ simulations was often lower (depending on the species), suggesting that SVs happening on regions with pairwise homology may be more difficult to resolve. As expected, the accuracy on ‘random’ SVs was higher than on more realistic simulations (‘known’ and ‘homologous’), suggesting that it may overestimate real data accuracy. In general, the F-value was higher than the ‘default’ setting in most species (except in *C. neoformans*), and the improvement was dramatic in some SV types (i.e. the F-value went from <0.1 to >0.95 in *C. glabrata*’s deletions or insertions) **(Figure 2)**. In addition, we found that parameter optimization increases recall rather than precision, which is >0.95 in most simulations and SV types **(Figure S3)**. Taken together, our results suggest that parameter optimization yields maximum performance by improving the recall of SVs as compared to default parameters.

**Figure 2.**
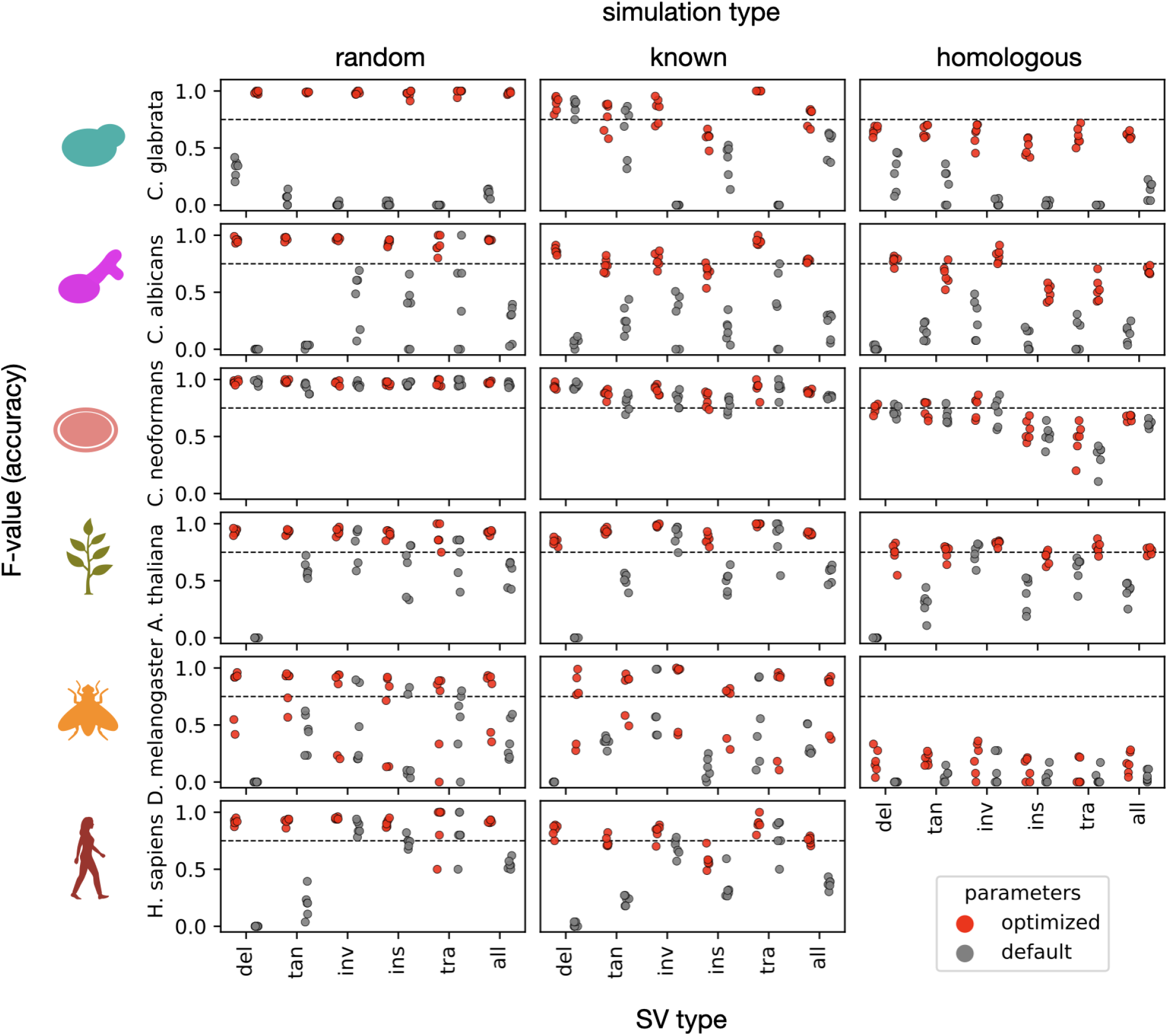
PerSVade’s parameter optimization improves the SV calling accuracy on simulations. We ran perSVade’s SV calling on three samples / species for six eukaryotes (see **Materials and Methods**) using either ‘random’, ‘known’ or ‘homologous’ simulations. These plots show the F-value of either default (grey) or optimized (red) parameters (for each sample and simulation type) on these simulations. The x axis represents the type of SV (deletions (del), tandem duplications (tan), inversions (inv), insertions (ins), translocations (tra) and the average of all SVs (all)). Note that **Figure S3** shows the corresponding precision and recall, from which the F-value is calculated.

We next explored whether different runs of perSVade (i.e. in different species or simulation types) yield similar parameters, which may clarify how necessary this optimization is. We hypothesized that each sample and simulation type combination may require specific parameters that would not necessarily work for other samples. To test this we first compared the chosen parameters across different runs, which appeared to be sample-specific **(Figure S4)**. This suggests that there is not a universal recipe (i.e. filtering parameters) for SV calling with perSVade. However, another (null) hypothesis could be that different parameter sets have similar outcomes, without changing the SV calling accuracy. This question was highly important to us. If perSVade’s optimization converges to equivalent parameter sets in different samples we would not need the optimization on every sample (i.e. we could re-define one of these parameters as default). In order to sort this out, we evaluated how different parameter sets (either ‘default’ ones or those that are defined as ‘optimum’ for a given sample) work on simulated genomes related to other samples. The results of this analysis are shown in **Figure 3** and **Figure S5**. As hypothesized, not all the parameter sets yield accurate results on all samples, with large differences between species **(Figure 3A)**. However, we found that parameters optimized for one sample are mostly accurate on samples of the same species, regardless of the simulation type **(Figure 3B)**. Of note, the parameters yielded by ‘random’ simulations were accurate on ‘homologous’ and ‘real’ simulations **(Figure 3)**. This indicates that running perSVade on ‘random’ simulations (the cheapest setting in terms of resources) yields accurate parameters for more realistic simulations and possibly real SVs. On another line, we found that the different parameters changed mostly the SV calling recall, and not the precision **(Figure S5)**. In summary, our results suggest that parameter optimization is necessary for maximum performance in each species and dataset.

**Figure 3.**
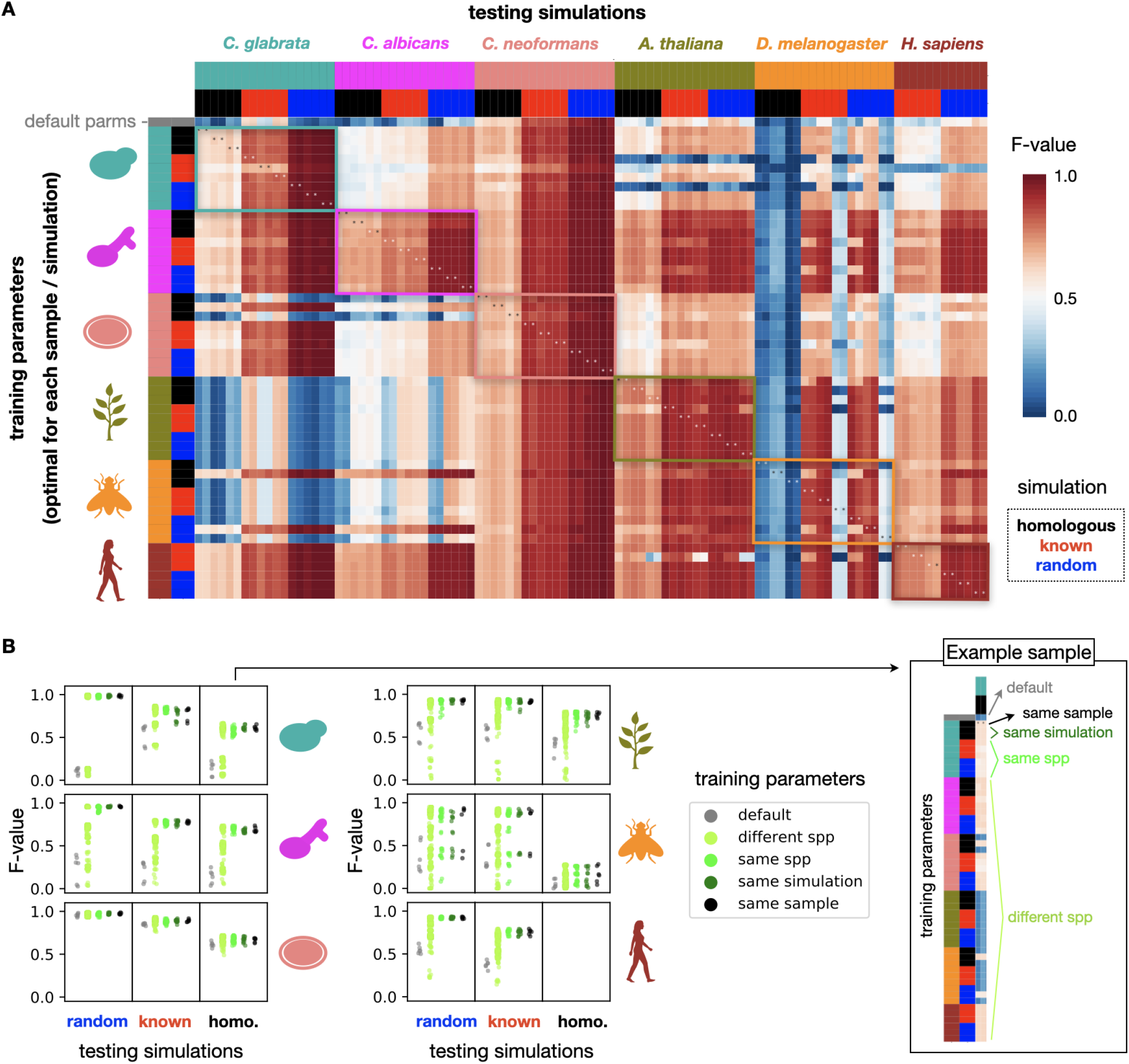
There is no universal recipe for SV calling across all species. **(A)** In order to assess whether perSVade’s parameter optimization is necessary for a given combination of sample and simulation (mentioned in **Figure 2**) we measured the SV calling accuracy of each optimized parameter set on the other combinations. Each row indicates a different “training” parameter set optimized for each sample and simulation type in all species. In addition, the first row refers to the default parameters. Each column represents a simulation from a given sample / simulation type to be “tested”. The heatmap shows the F-value of each parameter set on each tested simulation (hereafter referred to as ‘testing instance’). Note that the species are ordered alike in rows and columns. In addition, note that each sample (from a given species and simulation type) yielded one set of training parameters and two simulated genomes tested here, which explains why there are two columns for each row. The colored boxes indicate testing instances where the training and testing species are equal. The asterisks refer to instances where both the sample and type of simulation are equal in the training and testing (equivalent to the ‘optimized’ parameters from **Figure 2**). Note that **Figure S5** shows the corresponding precision and recall, from which the F-value is calculated. **(B)** We summarized the data shown in (A) to compare how similar types of training parameters performed on each species (in the rows) and type of simulations (in the columns). Each point corresponds to a testing instance, matching one cell from the heatmap in (A). The ‘default’ and ‘same sample’ reflect testing instances where the training parameters were either un-optimized or optimized specifically for each sample, respectively. The ‘different spp’ group includes instances where the training parameters were from different species. The ‘same spp’ group shows testing instances with both training parameters and tested simulations from a different sample of the same species. The ‘same simulation’ reflects instances with the same training and testing sample, but different simulation types. For clarity, the right box shows how the training parameters are grouped for a set of ‘homologous’ simulations based on one example *C. glabrata* sample (which corresponds to the first two columns in (A)).

### PerSVade’s parameter optimization improves the calling accuracy in datasets with defined sets of real SVs

The performance of SV calling on simulations may not be equivalent on real data, as SVs often appear around repetitive or low-complexity regions which hamper their detection (*24*, *34*–*36*). It is thus possible that we overestimated the real accuracy in our simulations. We partially addressed this with our analysis based on ‘realistic’ simulations (‘known’ and ‘homologous’), where the inferred accuracy was lower **(Figure 2)** and potentially closer to the real one. To further validate the usage of perSVade for real SV calling we tested it on datasets with known SVs, which were available for the human samples tested above (i.e. **Figure 3**). We ran perSVade (using different simulation types) on the same three datasets, which had previously-defined deletions and inversions (see **Materials and Methods** for details).

We used these data to assess the accuracy of perSVade on real datasets, using different sets of parameters (optimal for each simulation and sample from the six species tested above, shown in **Figure 3**). As expected, we found a lower F-value on real datasets **(Figure 4)** as compared to the simulated genomes **(Figure 2, 3)**, with high precision and lower recall **(Figure 4B)**. In addition, parameter optimization improved the F-value modulating both precision and recall **(Figure 4B)**. However, the other results described in the simulations’ analysis (related to the performance of the pipeline and the universality of the parameters) are qualitatively equivalent in these real datasets **(Figure 4)**. Taken together, our analysis indicates that perSVade improves SV calling in real datasets (similarly to simulated genomes).

**Figure 4.**
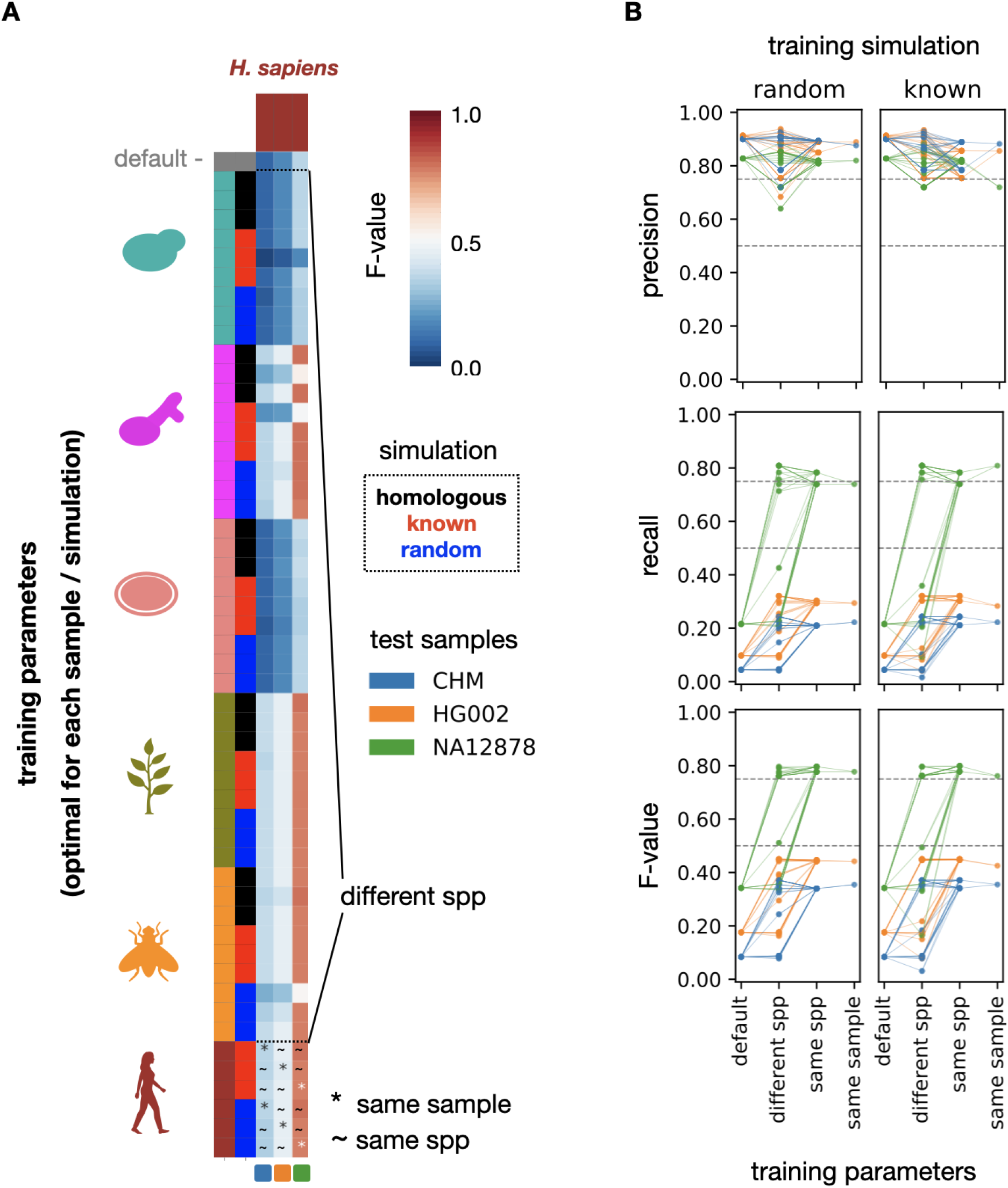
PerSVade’s parameter optimization improves the SV calling accuracy on datasets with known real SVs. **(A)** To test perSVade’s performance on real SVs we measured how the parameters optimized for several simulations in different species (see **Figure 3**) work on three human samples (CHM, HG002 and NA12878) with defined sets of real SVs. Each row indicates one of these different “training” parameters optimized for each sample and simulation type. In addition, the first row refers to the default parameters. Each column represents a sample with defined real SVs to be “tested”. The heatmap shows the F-value of each parameter set on each tested real sample (hereafter referred to as ‘testing instance’). In addition, we divide the testing instances into different groups (‘default’, ‘different spp’, ‘same spp’ and ‘same sample’), which are relevant to understand the (B) panel. The ‘different spp’ group refers to instances where the training and testing species were different. The ‘~’ (same spp) refers to instances where the training and testing samples were different, but from the same species. Finally, the ‘*’ (same sample) refers to instances where the training and testing samples were the same. **(B)** We summarized the data shown in (A) to compare how similar types of training parameters performed on each testing sample (each represented by a different color). Each row corresponds to a different accuracy measure. Each point corresponds to a testing instance (matching one cell from the heatmap in (A) in the bottom ‘F-value’ plots). The ‘default’ and ‘same sample’ reflect testing instances where the training parameters were either un-optimized or optimized specifically for each sample, respectively. The ‘different spp’ group includes instances where the training parameters were from a different, non-human, species. The ‘same spp’ group shows testing instances with both training parameters and tested simulations from different samples of the same species. In addition, each column represents testing instances where the training parameters were based on ‘random’ or ‘known’ simulations, respectively. Note that the different groups of ‘training parameters’ are equivalent to those shown in (A).

## DISCUSSION

Despite large variation of genomic features across taxa, SV detection approaches in non-model organisms tend to rely on tools and parameters developed for other species (generally human). We hypothesized that this “one size fits all” approach is suboptimal. To test this idea and overcome the problem, we developed perSVade, a flexible pipeline that automatizes the calling and filtering of structural variants (SV) across eukaryotes. PerSVade is a modular method to automatically adapt a state-of-the-art SV calling pipeline to any species of interest. PerSVade uses simulations to choose the optimal filters for each sample and report the calling accuracy, which can inform about the reliability of the results. This will allow users to be aware of the accuracy in their datasets (i.e. perSVade may be inaccurate in some datasets due to low coverage, short read lengths or excessive repeats in the genome) and make informed choices.

We validated the broad usability of perSVade by testing it on simulations and real datasets for a wide range of eukaryotes (with genomes of 12-3,000 Mb and datasets including 10^7^-10^9^ reads). We found that there is a significant computational burden related to parameter optimization, which may hinder its usage on large genomes. This means that perSVade may be particularly cost-effective for small genomes (i.e. <200 Mb), although the chosen settings will depend on the available resources.

This testing also revealed that, as we hypothesized, parameter optimization improves the calling accuracy on both simulations and datasets with real, previously-defined SVs. We found that the optimization mostly improves the recall rather than precision (which is generally high regardless of the used parameters). However, there are some exceptions (mostly in the testing on real SVs), suggesting that optimization can be necessary for reaching both high recall and precision in some samples. In addition, perSVade’s optimization yielded unique parameter sets for each sample, which were often inaccurate on other datasets. This means that there is no universal set of parameters that work well for all samples, which justifies the need for parameter optimization and a tool like perSVade to automate such a task. Conversely, we found some trends that can be useful to skip parameter optimization in some cases. For instance, parameter sets were often accurate across datasets of the same species. In addition, parameters resulting from ‘random’ simulations performed well in more realistic (‘known’ and ‘homologous’) simulations as well as in real SV datasets of the same species, indicating that they can be used for maximum performance. Based on these findings, we propose the following recommendations for a cost-effective usage of perSVade:

- For SV calling on many datasets of one species with similar properties (similar coverage, read length and insert size), run perSVade using ‘random’ simulations on one sample, and use the optimized parameters for the other samples (skipping optimization). The reported calling accuracy may be overestimated since the simulations are not realistic, but the chosen parameters are expected to be optimal.
- For approximating the real SV calling accuracy, run perSVade on realistic simulations (‘homologous’ or ‘known’), which may report an accuracy that is closer to the real one.

We note that perSVade is not a fundamentally new algorithm for SV detection but rather a pipeline implementing existing algorithms. This is why we did not compare it with other such methods (like *manta* (*20*) or *delly* (*62*)). The novelty of our pipeline lies in the automatic parameter selection feature, which is unique (to the best of our knowledge) for short read-based SV calling. We thus centered our testing on the accuracy of different parameters on SV calling. In fact, some recent approaches specifically developed for human genomes (*22*, *63*) may outcompete perSVade in human samples. However, such methods rely on previously-defined sets of known SVs, which are not available in most taxa. We thus consider that our pipeline will be mostly useful in species without such specific methods available. For example, perSVade was used in a recent study to find SVs associated with antifungal drug resistance in the non-model yeast *Candida glabrata* (*9*), which successfully validated all (8/8) the predicted variants using PCR.

Finally, perSVade also includes modules for CNV identification and SNP/INDEL calling, as a way to automate the finding of other broadly used genomic variants. In addition, it includes variant annotation features to ease the functional interpretation of these variants for downstream analyses. In summary, perSVade is a swiss-knife-like framework to identify many types of variants with a few bash commands. We consider that this tool will be useful to understand the role SVs in different phenotypes and organisms, particularly those with no specific recommendations.

## DATA AVAILABILITY

PerSVade is available at https://github.com/Gabaldonlab/perSVade and can be installed using either conda environments or through a docker image containing the pipeline, available at https://hub.docker.com/r/mikischikora/persvade. The github repository contains detailed examples on how to install and run perSVade using conda, docker or singularity. We have tested perSVade on several Linux and Mac architectures, and the docker image may be run in any machine in a reproducible way. All the results shown in this paper were generated using the script https://github.com/Gabaldonlab/perSVade/blob/master/scripts/perSVade.py from version 1.0, which is a wrapper to execute several modules with a single command. Since perSVade is an actively used (and maintained) pipeline, we have created a few new releases since version 1.0, which include an improved documentation, more unit tests and the implementation of an efficient debugging of inputs. Note that these changes do not affect the functionality of the modules as implemented in version 1.0. Hence, we recommend the usage of the latest version, which is the one with the best documentation and usability. All the data used for testing perSVade was obtained from the SRA database or public ftp servers, and is listed in **Table S1** and **Materials and Methods**. All the code necessary to reproduce the results and plots shown in this paper is in https://github.com/Gabaldonlab/perSVade/tree/master/testing.

## ACKNOWLEDGEMENTS

The authors thank Cinta Pegueroles and Marina Lleal for the useful discussions key in the building of perSVade. In addition, we want to thank Hrant Hovhannisyan, Valentina del Olmo, Diego Fuentes, Anna Vlasova, Maria Artigues, Matteo Schiavinato and Marina Marcet for beta-testing the pipeline and providing us with useful feedback.

## AUTHOR CONTRIBUTIONS

MAST wrote the code and performed all bioinformatic analysis. MAST and TG conceived the study, interpreted the results, and wrote the manuscript. TG supervised the project and provided resources.

