## Supplementary material for "PerSVade: Personalized Structural Variation detection in your species of interest"

### ADDITIONAL MATERIALS AND METHODS

#### Filters used by perSVade

These are the filters used in the module 'call_SVs', whose values may vary across parameter optimization in perSVade (note that most of the *gridss* filters were inspired by (*1*)):

- min_Nfragments: Minimum number of reads supporting a breakend in *gridss* to be accepted (default is 5).

- min_af: Minimum Variant Allele Frequency of a breakend in *gridss* to be accepted (default is 0.25).

- min_QUAL: Minimum quality (QUAL field of the vcf file) of a breakend in *gridss* to be accepted (default is 0).

- max_to_be_considered_small_event: Maximum length of a breakpoint in *gridss* to be considered a small event (default is 1000). Events shorter than this value are considered as "small events", which are treated particularly by other filtering steps.

- min_length_inversions: Minimum length of inversion-like breakends in *gridss* to be accepted (default is 40).

- maximum_lenght_inexactHomology: Maximum length of the inexact homology region around a breakend in *gridss* to be accepted (default is 50). This filter is not applied to "small events", as defined by "max_to_be_considered_small_event".

- maximum_microhomology: Maximum length of the exact homology (microhomology) region around a breakend in *gridss* to be accepted (default is 50).

- maximum_strand_bias: Maximum strand bias of a breakend in *gridss* to be accepted (default is 0.99). This filter is only applied to "small events", as defined by "max_to_be_considered_small_event".

- filter_noReadPairs: Discards *gridss* breakends without discordant read pair support (default is false). This filter is not applied to "small events", as defined by "max_to_be_considered_small_event".

- filter_noSplitReads: Discards *gridss* breakends without split-read evidence (default is false). This filter is only applied to "small events", as defined by "max_to_be_considered_small_event".

- filter_overlappingRepeats: Discards *gridss* breakends overlapping repetitive elements (default is false). This will only have an effect if you provide a repeats file as inferred by the module 'infer_repeats'.

- filter_polyGC: Discards *gridss* breakends with long inserted G or C sequences (>15bp) (default is true).

- wrong_FILTERtags: A set of values in the FILTER field of the *gridss* vcf which flag discarded breakends (default is ["NO_ASSEMBLY"]).

- range_filt_DEL_breakpoints: A range of lengths in which DEL-like breakends (as defined by *gridss*) are discarded if the breakend has a region with inexact homology above 5bp (default is [0, 1]). For example, if set to [500, 1000], DEL-like breakends whose length is between 500 and 1000bp with an inexact homology sequence >5 bp would be discarded.

- dif_between_insert_and_del: The margin given for comparing the length of the inserted sequence (len_seq) on a *gridss* DEL-like breakend and the length of the actual event (len_event) (default is 5). If len_seq > (len_event - dif_between_insert_and_del), the breakend is filtered out. This filter is only applied to "small events", as defined by "max_to_be_considered_small_event".

- max_rel_coverage_to_consider_del: The maximum relative coverage that a region spanning a DEL-like breakpoint (as defined by *clove*) can have to be classified as an actual deletion (default is 0.1).

- min_rel_coverage_to_consider_dup: The minimum relative coverage that a region spanning a TAN-like breakpoint (as defined by *clove*) can have to be classified as an actual tandem duplication (default is 1.8).

Note that all the breakpoints that have at least one breakend that does not pass the filters are discarded by perSVade.

### Calling of small variants

PerSVade's small variant calling pipeline (module 'call_small_variants') uses three alternative methods (GATK Haplotype Caller (HC) (*2*) (v4.1.2), freebayes (FB) (*3*) (v1.3.1) and / or bcftools (BT) (https://github.com/samtools/bcftools, v1.9)) to call and filter Single Nucleotide Polymorphisms (SNP) and small insertions/deletions (IN/DEL) in haploid or diploid configuration (specified with the *-p* option). The input is the .bam file generated by *bwa mem* (http://bio-bwa.sourceforge.net/bwa.shtml, v0.7.17) with the 'align_reads' module. This module defines as high-confidence (PASS) variants those that are in positions with a read depth above the value provided with *--min_coverage*, with extra filters for HC and FB. For HC, it keeps as PASS variants those where 1) there are <4 additional variants within 20 bases; 2) the mapping quality is >40; 3) the confidence based on depth is >2; 4) the phred-scaled p-value is <60; 5) the MQRankSum is >-12.5 and 6) the ReadPosRankSum is > -8. For FB, perSVade 'call_small_variants' keeps as PASS variants those where 1) quality is > 1 or alternate allele observation count is > 10; 2) strand balance probability of the alternate is > 0; 3) number of observations in the reverse strand is > 0; and 4) number of reads placed to the right/left of the allele are > 1. Then, bcftools (v1.10) and custom python code are used to normalise and merge the variants called by each software into a consensus variant set, which includes only those variants called with high-confidence by *N* or more algorithms This results in one .vcf file with the high-confidence variants for each *N*. Note that this .vcf file only keeps variants for which the fraction of reads covering the alternative allele is above the value provided with *--min_AF* (which may be 0.9 for haploids or 0.25 for diploids). For diploid calls, it defines the genotype with the strongest support (the one called by most programs). In addition, the quality of each variant is calculated from the mean of the three algorithms. Beyond the filtered variant calls, this module writes a tabular file with all the raw variants with various metadata columns (i.e. the programs that called the variant), which can be used to apply a custom filtering of the variants.

### Comparing sets of SVs to calculate precision and recall

To measure accuracy in different sets of "called SVs" (in perSVade's simulations and also the testing of the pipeline (related to **Figures 2, 3, 4, S1, S3, S5**)) we compared them against the corresponding sets of "known SVs" and calculated the following estimates:

- precision = TP / (TP+FP)
- recall = TP / (TP+FN)
- F-value = (2 * precision * recall) / (precision + recall)

Where true positives (TP) are those in the “called SVs” that match at least one variant from the “known SVs”, false positives (FP) are those in the “called SVs” that do not match any from the “known SVs” and false negatives (FN) are those in the “known SVs” that are not matched by any variant from the “called SVs”. We define that two SVs are “matching” using a different approach for each type of SV:

- Inversions, tandem duplications and deletions: both SVs are in the same chromosome, their altered regions are overlapping by 75% and their breakends are <50bp apart.

- Insertions: both SVs have the same origin and destination chromosomes and are both either cut-and-paste or copy-and-paste. In addition, the regions of the origin chromosome are overlapping by 75% and the breakends are <50bp apart. Finally, the starts of the destination chromosomes (insertion sites) in both SVs are <50bp apart.

- Translocations: both SVs have the same origin and destination chromosomes and are both either inverted or not. In addition, the breakpoint positions in both SVs are <50bp apart.

In addition, we calculated ‘integrated’ precision and recall measures (related to **Figures 3, 4 and S5**)) merging all the variants together into single sets of “called SVs” and “known SVs”. We used custom python (v3.6) code and *bedmap* from the *bedops* tool (*4*) (v2.4.39) to calculate all these overlaps. See **Materials and Methods** for further information on the meaning of each type of SV.

### SUPPLEMENTARY FIGURES

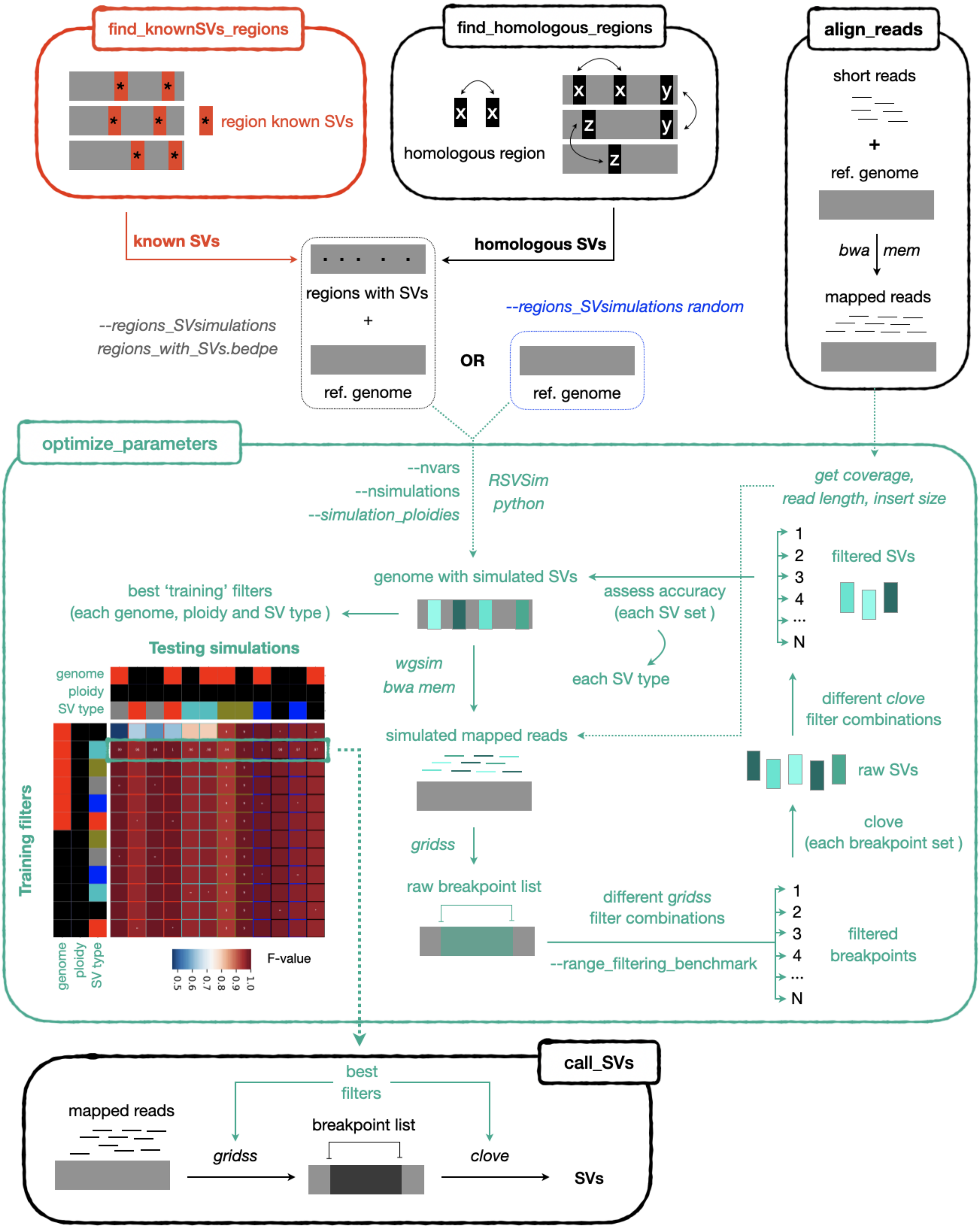


**Figure S1. Detailed workflow of the 'optimize_parameters' module.** The module 'optimize_parameters' is the core, most novel function of perSVade. It requires the argument --regions_SVsimulations, which specifies the regions of the genome for simulations of SVs. These can be either around some specific regions (with the argument --regions_SVsimulations <regions>.bedpe) or randomly placed across the genome (with the argument --regions_SVsimulations random). Note that perSVade has modules to infer either regions with previously known SVs (through 'find_knownSVs_regions') or regions with pairwise homology (through 'find_homologous_regions'). The SV simulations around such regions may be more realistic than random simulations, which is why they may be considered. The module 'optimize_parameters' finds a set of optimum parameters through simulations around these regions. By default, it generates two simulated genomes (tunable with --nsimulations) with 50 SVs of each type (tunable through --nvars) based on the reference genome and the provided regions. There are two simulated genomes for each of the desired ploidies, tunable through --simulation_plodies. For example, we set '--simulation_plodies haploid' for haploid organisms and '--simulation_plodies diploid_hetero' for diploids (which means that the simulated genomes will have only heterozygous variants) in the testing of perSVade on several organisms (see **Materials and Methods**). For each simulated genome perSVade 'optimize_parameters' simulates reads with equal insert size, coverage and read length as the input mapped reads (provided with the argument -sbam). Then it aligns the reads and runs gridss to obtain a list of 'raw breakpoints'. This module then tries several combinations of filters on them (by default >278,000,000, which is tunable through --range_filtering_benchmark) to generate many 'filtered breakpoints'. Each of these is processed with clove to generate a set of 'raw SVs'. PerSVade 'optimize_parameters' next tries several combinations of filters on each of them to get a set of filtered SVs. These are compared against the true set of SVs (inserted in the simulated genome) to calculate the accuracy (F-value) of each combination of gridss and clove filters on each simulated genome, ploidy and SV type. These filters are optimised for each simulation, and thus may not be accurate on independent sets of SVs (due to overfitting). In order to reduce this effect, this module tests how each of these 'best' filters perform on all simulations, ploidies and SV types (not only in those that yielded the given filters as optimum). The heatmap shows the F-value for an example sample (BG2 based on random simulations from *Candida glabrata,* see **Materials and Methods**), where the filters in the second row are accurate on all simulations (indicating that there is no overfitting on them) and thus they are chosen as the final set of best parameters. Note that the filters in the first row are only accurate on some simulations, suggesting that they are overfitted and thus they are not chosen as good filters. At the end, this module writes the accuracy of these best parameters into a .tab file, which will allow the user to understand how much the results are to be trusted. In addition, these optimised filters (or parameters) are written into a .json file that may be used for calling SVs with 'call_SVs'.

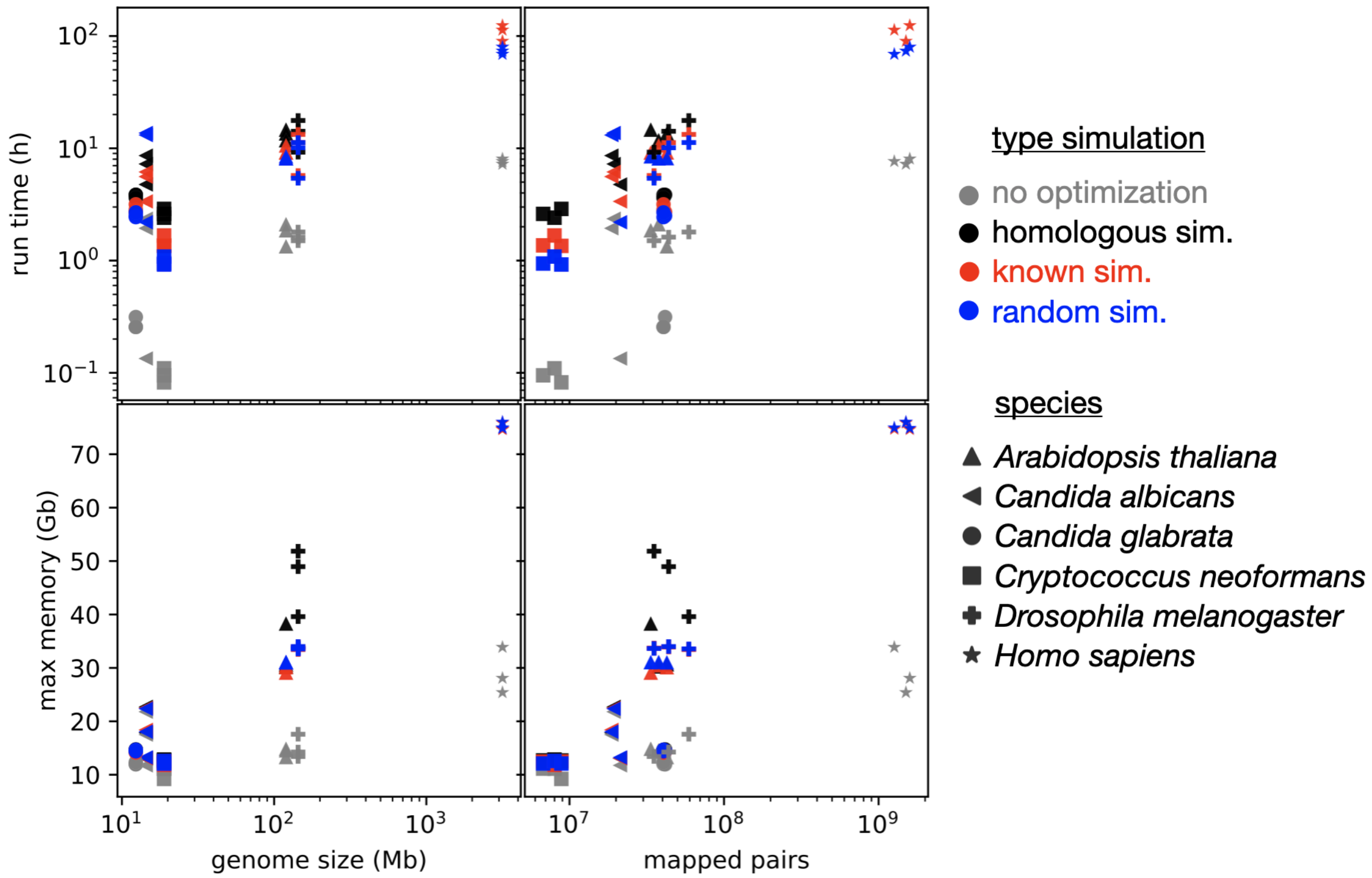


**Figure S2. PerSVade's parameter optimization requires extra resources.** We tested perSVade's SV calling modules ('optimize_parameters', 'call_SVs' and 'integrate_SV_CNV_calls') on six eukaryotes (three samples per species) using either no parameter optimization (gray) or different types of simulations (black, red, blue) for the 'optimize_parameters' module in a machine with 16 cores. Shown are the running time and maximum RAM used ignoring the resources related to read alignment (which was performed independently). Thus, each point reflects the resources used by 'optimize_parameters' (except in the gray points), 'call_SVs' and 'integrate_SV_CNV_calls'. Of note, perSVade was run with a different setting for the human datasets to avoid excessive resource consumption. First, we skipped the marking of duplicate reads on the .bam files. Second, we ran the simulations on a subset of the genome (only chromosomes 2, 7, 9, X, Y and mitochondrial). Third, we skipped the 'homologous' simulations in human samples due to excessive memory consumption. The x axes reflect the reference genome size (left) and the number of mapped read pairs (right), which are correlated with resource consumption. This data may be useful to allocate computational resources for running perSVade.

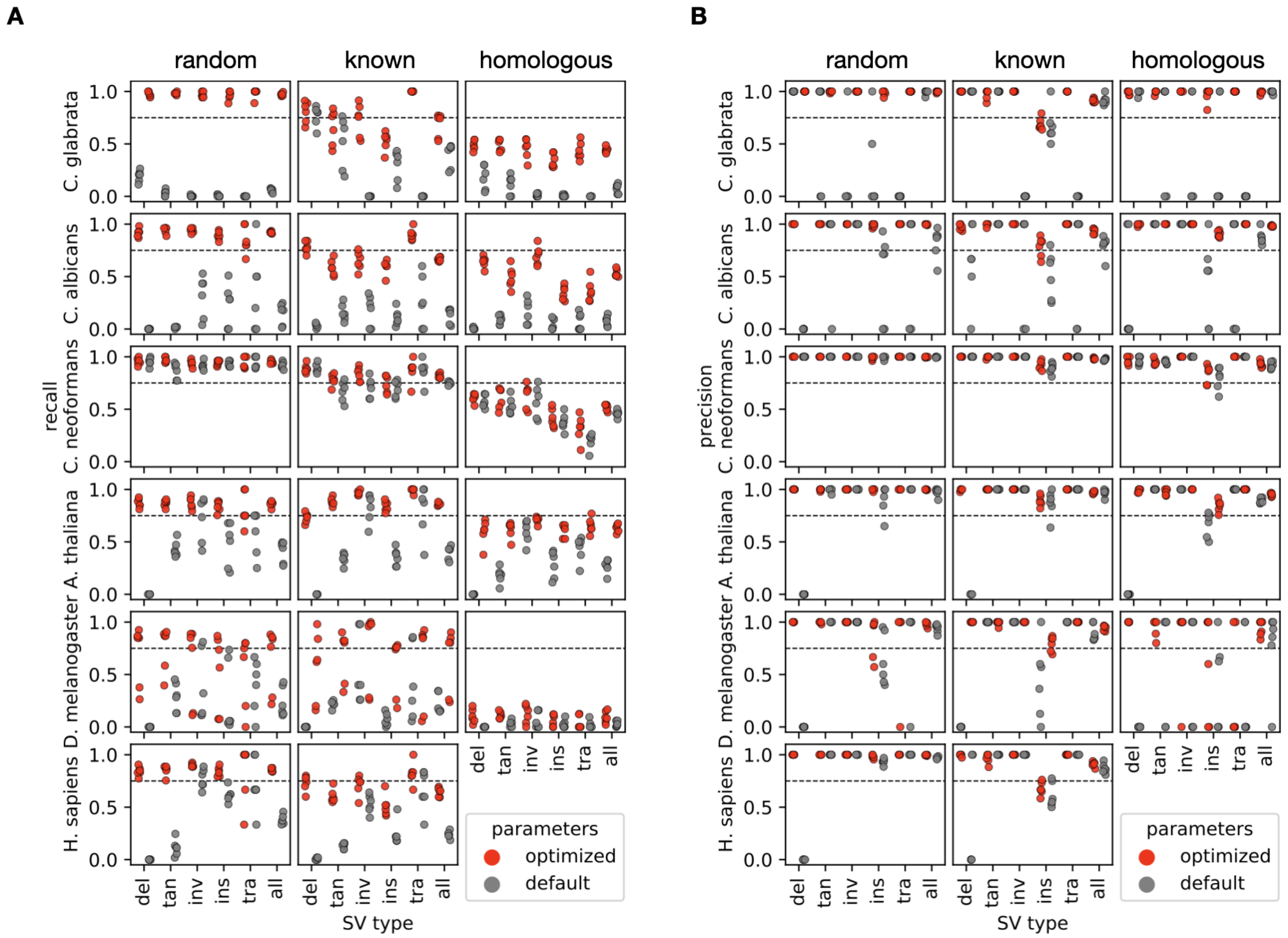


**Figure S3. PerSVade's parameter optimization improves the recall of SVs.** We ran perSVade's SV calling modules on three samples per species for six eukaryotes (see **Materials and Methods**) using either 'random', 'known' or 'homologous' simulations. These plots show the recall (left) and precision (right) of either default or optimized parameters (for each sample and simulation type) on these simulations. The x axis represents the type of SV (deletions (del), tandem duplications (tan), inversions (inv), insertions (ins), translocations (tra) and the average of all SVs (all)).

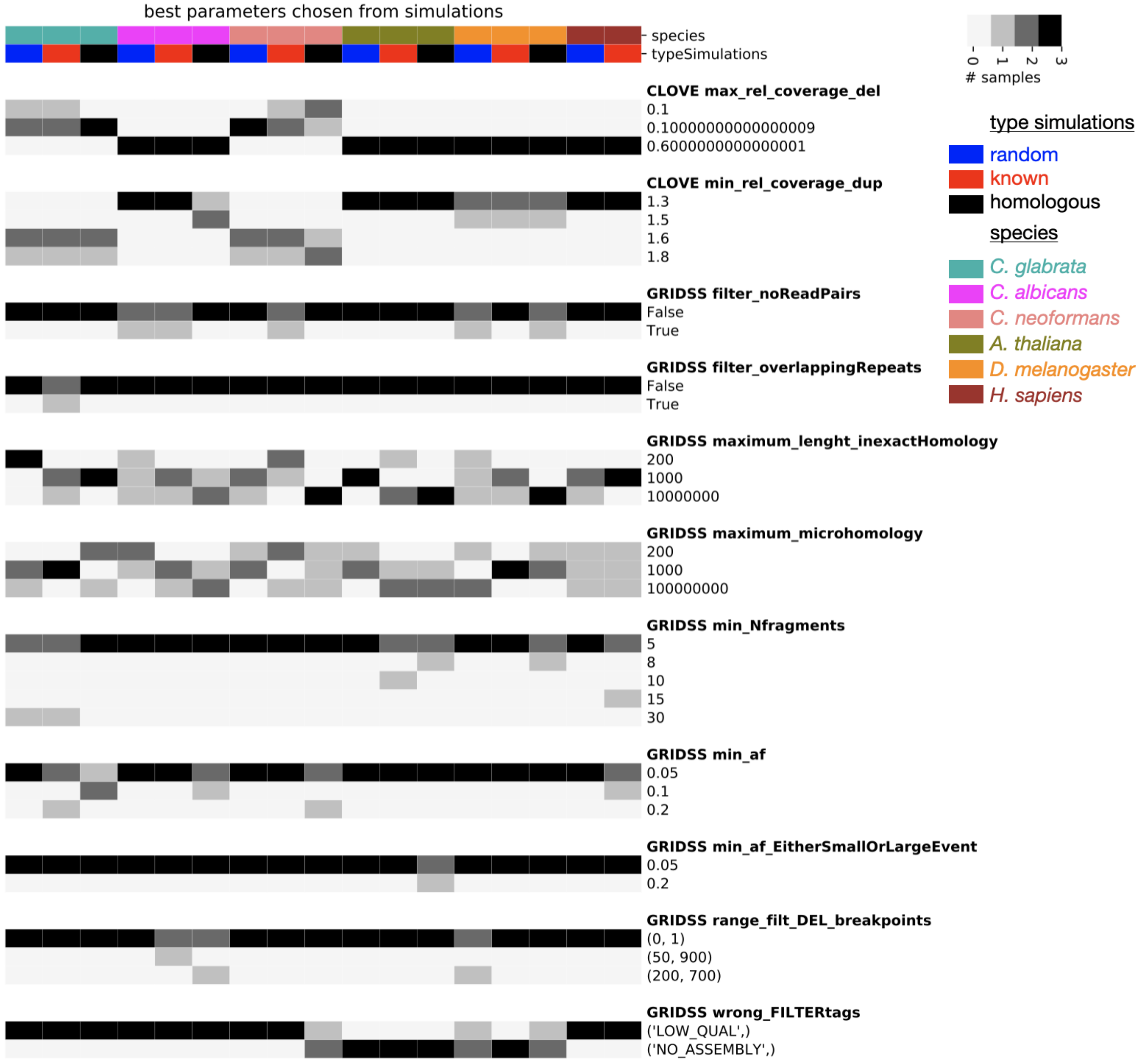


**Figure S4. Each sample yields a different set of optimum parameters.** We ran perSVade's 'optimize_parameters' module on six eukaryotes (three samples per species) and a parameter optimization based on either 'random', 'homologous' or 'known' simulations (see **Materials and Methods**). Shown are the chosen values for each parameter (only those that changed across samples) in each optimization procedure. Each group of rows refers to the values chosen for one type of parameter (see **Additional Materials and Methods**) used for *gridss* or *clove*. The color indicates how many samples (from zero to three) yielded a specific value for each parameter type. For example, the threshold to discard breakpoints (called by *gridss*) based on allele frequency was set to 0.05 in most samples (see "GRIDSS min_af"). However, the optimization for 'known' SVs in *C. glabrata* yielded a threshold of 0.2 in one sample (out of three) (see the second column).

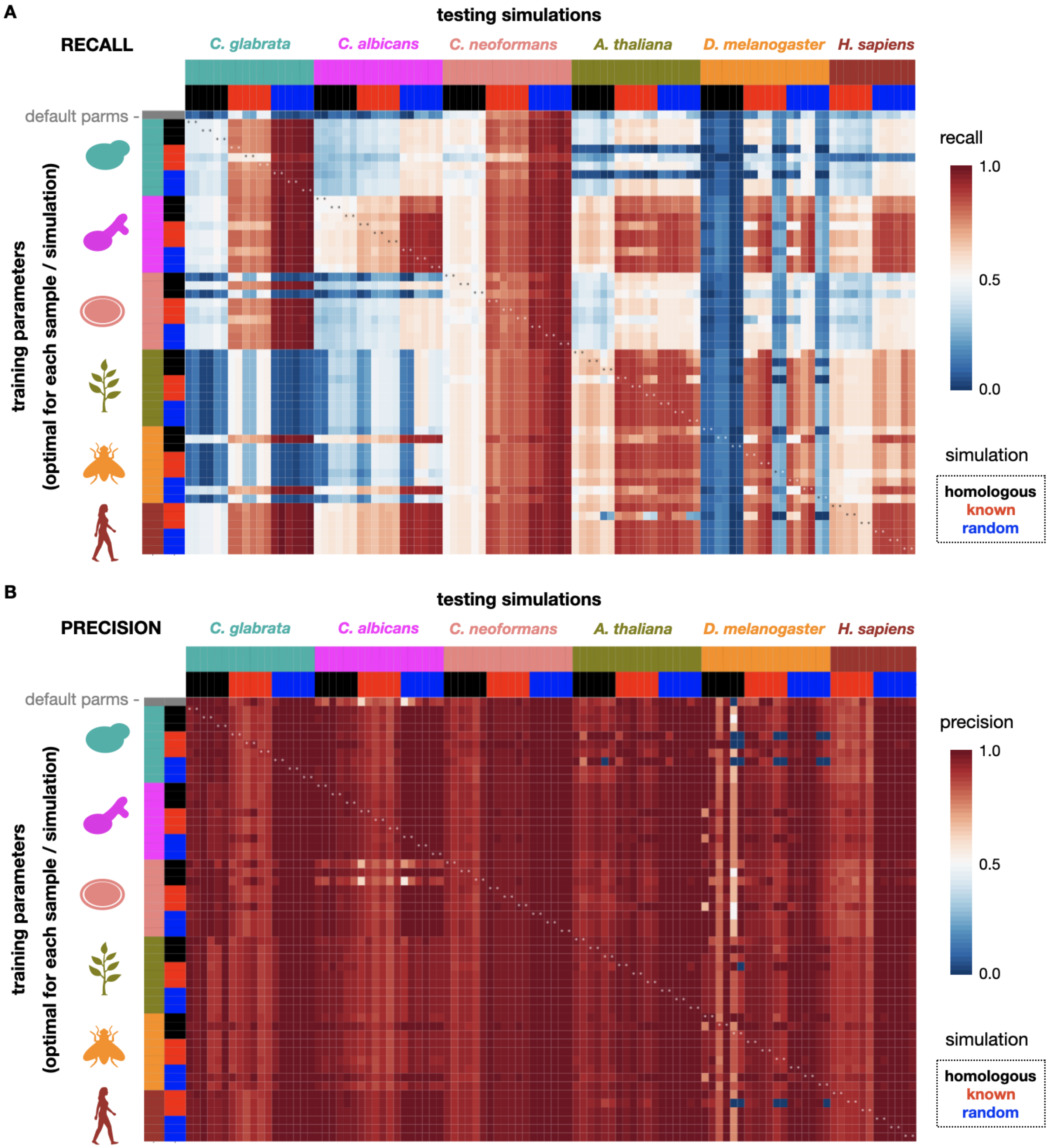


**Figure S5. PerSVade's parameters optimization mostly changes the recall of SVs in simulations.** To assess whether perSVade's parameter optimization is necessary for all samples / simulations (mentioned in **Figure 2 and S3**) we measured the SV calling accuracy of each parameter set on the other samples / simulations. Each row indicates a different "training" parameter set optimized for each sample and simulation type in all tested species. In addition, the first row refers to the default parameters. Each column represents a simulation from a given sample / simulation type to be tested. The heatmap shows either the recall (A) or the precision (B) of each parameter set on each tested simulation. Note that the species are ordered alike in rows and columns. In addition, note that each sample (from a given species and simulation type) yielded one set of "training" parameters and two simulated genomes tested here, which explains why there are two columns for each row. The asterisks refer to testing instances where both the sample and type of simulation are equal in the training and testing (equivalent to the 'optimized' parameters from **Figure 2 and S3**)

### SUPPLEMENTARY TABLES

| **target_species** | **target_taxID** | **sample_taxID** | **sample_species** | **SRA_run** | **% reads map.** |
|---|---|---|---|---|---|
| *C. glabrata* | N/A | N/A | *C. glabrata BG2* | SRR15498429 | N/A |
| *C. glabrata* | N/A | N/A | *C. glabrata* CST34 | SRR15498440 | N/A |
| *C. glabrata* | N/A | N/A | *C. glabrata M12* | SRR15498481 | N/A |
| *C. albicans* | 5476 | 1182531 | *C. albicans 3153* | SRR641729 | 94.58 |
| *C. albicans* | 5476 | 1182537 | *C. albicans A123* | SRR538772 | 88.77 |
| *C. albicans* | 5476 | 1182540 | *C. albicans A203* | SRR538786 | 95.91 |
| *C. neoformans* | 5207 | 1423894 | *C. neoformans Bt35* | SRR1063293 | 99.93 |
| *C. neoformans* | 5207 | 1423915 | *C. neoformans RSA-MW-1281* | SRR1063017 | 99.86 |
| *C. neoformans* | 5207 | 1423916 | *C. neoformans RSA-MW-5465* | SRR1063214 | 99.94 |
| *A. thaliana* | 3702 | 38785 | *A. arenosa* | SRR4128971 | 76.76 |
| *A. thaliana* | 3702 | 378006 | *A. arenosa x A. thaliana* | ERR5032500 | 89.59 |
| *A. thaliana* | 3702 | 2608267 | *A. arenosa x A. lyrata* | ERR3514861 | 65.98 |
| *D. melanogaster* | 7227 | 7238 | *D. sechellia* | SRR5860659 | 89.70 |
| *D. melanogaster* | 7227 | 7240 | *D. simulans* | ERR1597900 | 84.44 |
| *D. melanogaster* | 7227 | 7243 | *D. teissieri* | SRR13202235 | 86.61 |

**Table S1. Datasets used for the testing in simulations in C. glabrata, *C. albicans, C. neoformans, A. thaliana* and *D. melanogaster*.** We chose these datasets automatically from the SRA database for *C. albicans, C. neoformans, A. thaliana* and *D. melanogaster*. In order to have enough SV calls we selected mildly divergent samples (as compared to the reference genome) with a NCBI taxonomy taxon ID (indicated by each sample_taxID) different from the ID of species of interest (target_taxID). However, we only kept samples with most reads mapped (specified in the column '% reads map.') in order to discard datasets from highly divergent taxa. Note that it was not possible to find such samples for *C. glabrata* at the time of this study. We thus used three datasets for *C. glabrata* strains from our lab (*5*). 'N/A' indicates that the column (i.e. taxID or % of mapped reads) was not taken into consideration for selecting these samples. See **Materials and Methods** for more information.
